# Application of Ensemble Machine Learning to Metabolomic Data Identifies Metabolites Associated with Macrophage Polarization

**DOI:** 10.1101/2024.10.11.617846

**Authors:** Aadi Gannavaram, Rahul Paul, Spyros Karaiskos, Marissa Howard, Lance Liotta, Padmanabhan Seshaiyer

**Affiliations:** Montgomery Blair High School, Silver Spring MD USA; Division of Analytics & Benefit-Risk Assessment, Center for Biologics Evaluation and Research, US Food and Drug Administration, Silver Spring MD USA; Center for Applied Proteomics and Molecular Medicine, George Mason University, Fairfax VA; Mathematical Sciences Department, George Mason University, Fairfax VA

**Keywords:** Macrophage polarization, Metabolomics, AI/ML methods, ODE models, Topological data analysis

## Abstract

Towards developing quantitative models of anti-tumor activities of macrophages, we evaluated the effects of cytokines, tumor exosomes, and polarization states of macrophages in a tumor microenvironemnt using a system of differential equations. We modeled the non-linear dynamics of macrophage polarization states (M0/M1/M2), tumor cell killing by macrophages, and evasion of macrophage mediated killing by tumor originated extracellular vesicle decoys. Solving these coupled differential equations using numerical approaches, showed that the rate of macrophage polarization into the M1 state is the critical determinant of anti-tumor activity mediated by M1 polarized macrophages. To determine what metabolomic factors correlate with the polarization of naïve macrophage into anti-tumor M1 or pro-tumor M2 phenotypes, we performed LC/MS-based untargeted metabolomic analysis. Statistical analysis using Python-Scikit-learn was performed on the metabolomic data from naïve, M1 or M2 polarized murine macrophages followed by multiple feature selection methods. Application of ensemble machine learning methods to both secreted and cell associated metabolites revealed novel molecules of fatty acid metabolism to be the main mediators of polarization. Integration of ensemble machine learning feature-ranking tools into our analysis of metabolomic data identified new potential targets in macrophage metabolism for enhancing anti-tumor activities.

## 1. Introduction

The tumor microenvironment (TME) is a biological ecosystem comprising a variety of cell types [1]. TME is a complex network of interactions between tumor cells, immune cells, non-cellular components such as the extracellular matrix, exosomes, and secreted molecules such as cytokines and chemokines [2]. Depending on the milieu, immune cells such as macrophages may acquire the capacity to kill off tumor cells. The tumor cells, in contrast, employ various evasion mechanisms to neutralize the anti-tumor activities. Due to the high energy requirement of the rapidly proliferating tumor cells, the tumor cell metabolism can affect the TME to limit immune responses, and diminish the effectiveness of cancer therapies [3]. The expanding field of immune metabolism showed novel avenues to reshape the TME to promote anti-cancer immunity by altering cell metabolism [4]. Identifying targets that alter cancer metabolism towards bolstering inflammation will help augment the efficacy of cancer therapies.

Towards enhancing the effectiveness of therapies, the interactions between tumor cells and immune cells such as macrophages are intensively studied [5]. Monocytes, the precursor cells, may differentiate into dendritic cells or macrophages that perform surveillance functions, recognize pathogens, tumor cells and exhibit phagocytic functions. Naïve (M0) macrophages in the TME can polarize towards a pro-inflammatory, classically activated M1 or alternatively activated, anti-inflammatory M2 phenotypes depending on the cytokine and metabolic milieu [6]. The M1 polarized macrophages display anti-microbial and anti-tumor activity and the M2 type display tumor-supportive activity in the TME [7].

*In vitro* differentiation of macrophage cultures facilitates the analysis of immune metabolism and is widely used to identify novel targets for therapy. Accordingly, naïve (M0) macrophages are polarized to a M1 phenotype by treatment with the bacterial cell product lipopolysaccharide (LPS) or Th1 cytokines such as IFN-gamma and TNF-alpha.

The M1 polarized macrophages are identified by the expression of cell surface markers MHC-II, CD80, CD86, and the capacity to produce anti-tumor/anti-microbial iNOS,, pro-inflammatory cytokines including TNF-alpha, IL-1beta, IL-6, and chemokines such as CXCL9. [8]. The cytokines/chemokine signals produced by M1 polarized macrophages in turn polarize naïve macrophages to M1 phenotype. Transcription factors such as STAT1 and NF-κB are the key mediators of pro-inflammatory activity in M1 macrophages in anti-microbial and anti-tumor contexts [9]. In contrast, M2 polarization is induced by Th2 cytokines including IL-4, IL-13, IL-10, and TGF-beta. M2 macrophages characteristically express cell surface markers such as CD206, CD163, and FIZZ1. Similar to M1 macrophages, M2 macrophages attract naive macrophages and polarize them to the M2 phenotype by secretion of anti-inflammatory cytokines and chemokines including IL-10. TGF-beta, CCL17, and CCL22 [10].

The anti-tumor activities in the TME by the M1 macrophages can be affected through direct cytotoxic activity, such as reactive oxygen species (ROS), or secretion of IL-1beta and TNF-alpha. M1 macrophages also engage in indirect cytotoxic activity through the recruitment of NK cells and T cells that recognize tumor cells and induce apoptosis [11]. In contrast, M2 macrophages support tumor cell growth by secreting adrenomedullin and vascular epithelial growth factors (VEGFs) and releasing immunosuppressive molecules including IL-10, PD-L1, and TGF-beta [12, 13].

The relationship between metabolism of macrophages and their biological functions has become very clear [14]. The metabolic states of the macrophage subsets underpin their functional characteristics [15]. For instance, M1 macrophages mainly rely on aerobic glycolysis since this pathway can provide energy needed for microbicidal and tumoricidal activity rapidly [15]. This involves greater conversion of pyruvate to lactate and attenuation of respiratory metabolic pathways such as the tricarboxylic acid (TCA) cycle, allowing for higher reactive oxygen species (ROS) production. The pentose phosphate pathway (PPP) is also induced in M1 macrophages, generating NADPH for NADPH oxidase, which is necessary for ROS production and nitric oxide synthesis [16]. Additionally, several proinflammatory cytokines also induce the activation of NF-κB, a transcriptional regulator of macrophage function, which upregulates the expression of genes including HIF1α. This results in the upregulation of glycolytic enzymes such as GLUT1, which increases glucose intake, and lactate dehydrogenase, which produces lactate from pyruvate and prevents the latter from entering the M2-associated Krebs cycle [17, 18].

M2 macrophages, in contrast, rely heavily on oxidative metabolism. M2 macrophages are characterized by increased supply of pyruvate to the Krebs cycle, attenuation of the PPP, and a fully intact TCA cycle that allows long-term ATP generation [15]. M1 macrophages are characterized by a truncated TCA cycle after citrate and succinate production, leading to the accumulation of metabolites such as citrate, itaconate, and succinate [19]. Thus, creating a glycolytic environment can promote M1 polarization. For example, switching isoforms of the PFK2 enzyme to one that maintains higher fructose-2,6-bisphosphate concentration increases glycolytic flux and the expression of inflammatory cytokines. This mechanism is independent of traditional HIF1α expression, as even when the gene expression is inhibited, the PFK2 switch still increases glycolytic flux in classically activated (M1) macrophages [20].

Current efforts to model macrophage polarization in the TME mainly focus on the cellular-level interactions, such as systems of differential equations or agent-based modeling in a spatio-temporal platform [21-23]. These models focus on cytokine/chemokine-based interaction between local TME cell populations to identify important factors and effects of macrophage polarization in the TME. Given the essential role of metabolism in dictating the polarization of macrophages, it is important to develop models of the interactions in TME to understand the quantitative relationships. Additionally, liquid chromatography–mass spectrometry (LC–MS) based methods are widely used to obtain metabolomic data for investigations on anti-cancer therapies. Statistical methods and web-based tools for annotation of features, identification of perturbed pathways exist, however, machine learning methods are not widely applied for the analysis of metabolomic data sets [24]. Application of ML methods to metabolomic data from differentially polarized macrophages may help identify new targets for alteration towards enhancing therapies against cancer.

Mathematical modeling, analysis, and simulation using differential equations is quickly becoming the foundation of approaches used to solve multidisciplinary problems in science, engineering, and medicine [25, 26]. In the past, ordinary differential equation (ODE) models have been introduced for intracellular cytokine-metabolite interactions but focus on the impact of cytokines on changes in metabolism rather than assessing the effect of particular metabolites on polarization, and identifying the most critical metabolites [27]. To determine the most determinant metabolites in macrophage polarization, ensemble-based genome-scale modeling incorporates transcriptomics data with existing gene expression and metabolomic pathway methods (such as iMAT, GIMME, CORDA, INIT, and FASTCORE) to create a network model of metabolomic pathways involved in macrophage polarization. Then, a flux-weighted PageRank algorithm is implemented to determine the most important pathways by relative importance [28].

In this study, we analyzed the interactions of tumor cells, macrophages and the exosomes that result in the polarization of macrophages towards either tumor promoting or anti-tumor phenotypes. Modeling of these activities based on the observed parameters through ordinary differential equations allowed us to isolate key parameters that influence the anti-tumor activities of macrophages. Since the functional states of the macrophages are determined by the metabolism, we performed untargeted LC-MS analysis of in vitro differentiated macrophages. Using metabolomic data, we applied ensemble-based feature selection methods to identify metabolites that are associated with macrophage polarization states. The ensemble-based feature selection methods employed in this study do not rely on any network model, and exclusively select the most important singular metabolites for targeting based on metabolite abundance data. This machine learning classifier-based approach could produce more accurate results due to it not incorporating any preexisting models of metabolic pathway interactions, instead only taking into account the role of each individual metabolite in determining macrophage polarization. Finally, we applied topological data analysis methods to validate the findings from the ensemble-based methods. The proposed methodology is particularly useful for small biological datasets to ensure resistance to overfitting/bias, and reproducibility of the results. The metabolites identified could be targeted for macrophage polarization in a potential cancer therapy.

## 2. Materials and Methods

Mathematical model: To develop a quantitative understanding of the relative contribution of the factors in macrophage polarization, a system of ordinary differential equations representing various parameters such as the rate of change in the populations of tumor cells, M0/M1/M2 macrophages, and the quantity of tumor-secreted EVs in the simulated tumor microenvironment is used based on the literature. The simluations were run using the equations and parameters found in the published literature.

Macrophage polarization: Murine macrophages (RAW 264.7) were cultured in 24 well plates at a density of 0.5×10^6^/ml in complete RPMI media. To differentiate naïve macrophages into M1 type, 20ng/ml IFN-γ and 1ng/ml LPS were added to cultured media for 24hrs. To obtain M2 type macrophages 20ng/ml IL-4 was added and cultured for 48 hrs [29]. Cell pellets and culture supernatants were collected separately for mass spectrometric analysis.

Quenching and extraction of adherent cells: Macrophages were washed with room temperature PBS three times to thoroughly remove any remaining culture medium. To quench ice-cold 80% (−48 °C) (v/v) methanol: water (1 mL per T75 flask) was added to cover all the cells. Cells were harvested using a disposable cell scraper to remove all the cellular monolayer from the flask surface followed by aspiration of the quenching solution. To extract the metabolites, the sample tube was submerged into liquid N2 for 30 s to snap freeze the cells and then allowed to thaw on dry ice. This process was repeated three times with 10 s vortex mixing between cycles. The solution was centrifuged at - 9 °C, 11,500 g for 10 min and supernatants were collected. Samples were lyophilized overnight (∼ 14 h) and the pellets were stored at -80 °C immediately.

Preparation of cell supernatant for metabolomics analysis: Cell supernatant was removed from the cultures and centrifuged at 13,500 g for 5 min to remove any cell debris and suspended cells. The supernatant was filtered filter using a 0.45 μm syringe filter. Samples containing 200 μL supernatant were lyophilized samples overnight (∼14 h) until dryness and stored at -80°C until further analysis.

Mass spectrometry: The metabolites extracted from the macrophage cell pellets (n=3 each from M0, M1 and M2 cultures) and culture supernatants (n=3 each from M0, M1 and M2 cultures) were subjected to untargeted LC-MS analysis performed on a Thermo Orbitrap LTQ XL with HPLC separation on a Poroshell 120 SB-C18 (2 × 100 mm, 2.7 μm particle size) with an WPS 3000 LC system. The gradient consisted of solvent A, H_2_O with 0.1 % Formic acid, and solvent B 100 % acetonitrile at a 200 μL/min flow rate with an initial 2 % solvent B with a linear ramp to 95 % B at 15 min, holding at 95% B for 1 minutes, and back to 2 % B from 16 min and equilibration of 2 % B until min 32.5 μL was injected for each sample and the top 5 ions were selected for data dependent analysis with a 15 second exclusion window.

Feature annotation: Annotation of metabolites from the data obtained from the untargeted LC-MS analysis, including database comparison and statistical processing was performed in Progenesis QI. The pooled sample runs were selected for feature alignment. We proceeded with database matching using the Human Metabolome Database, selecting for adducts M+H, M+Na, M+2H, and 2M+H for positive mode and M-H, M+Cl, M-2H, and 2M-H and less than 10 ppm mass error, unique features were tentatively identified as potential metabolites. Metabolites were only annotated with MS/MS fragmentation matching scores above 20% with Progenesis Metascope. Using the pooled QC samples, features with above 30% CV and max abundances below 5000 intensity were filtered out and ANOVA p-value scores between the groups were calculated with a cutoff of < 0.05.

ML Methods: To discover important metabolic biomarkers and to understand the polarization of naïve macrophage into anti-tumor M1 or pro-tumor M2 phenotypes, ML feature selection methods were used followed by classifiers (M1 vs. naïve or M2) to evaluate classification performance. The LC-MS data was used in feature selection, with the experimental replicates as samples and each annotated metabolite as a feature. The data set had 9 samples (rows) and 503 features (columns) with 1 additional column to indicate classification (Supplementary information). We utilized 3-fold cross validation (cv) to evaluate the classification performance. Following cv fold splitting, data scaling (MinMaxScaler) was performed separately on the training (6 samples) and test data (3 samples) in 30 iterations. The cv model’s training set was balanced using synthetic minority oversampling (SMOTE) prior to feature selection. For each of the 3 fold cv, feature selection was conducted using a leave one out cv method. Three different feature selection strategies were chosen for our study: ANOVA F-Score, chi-square and logistic regression (lasso) for an ensemble feature selection strategy. Nested cross validation combined with ensemble feature selection would reduce bias and produce stable biomarkers.

The use of multiple feature selection algorithms is prioritized over a single method/algorithm to ensure the robustness of the feature selection and resistance to overfitting for any decision model that will be designed based on the selected meaningful and representative features with respect to the classification task. Each feature selection method is weighted equally and the set of cumulative relevant features is determined in the following way:

1. Ranking the individual features over 30 runs for each feature selection algorithm
2. Combining the selected features from each algorithm into a single cumulative list
3. Selecting the top features most frequently selected features in the list for M1 vs non-M1 macrophage classifier (Random forest for resistance to overfitting selected)

Due to the nature of our dataset, we opted to use feature selection methods applicable but not limited to large datasets. Hence, we did not use random forests, boosting, or similar algorithms particularly suited for large datasets. The three feature selection methods were chosen due to their effectivness with low sample sizes. After the feature selection, **a** random forest model was built on the subset of total features. We chose to build a random forest classifier since they are resistant to overfitting. The model is a classifier of M1 macrophages and non-M1 (naive, M2) macrophages based on the metabolomic profile. The random forest models are used as a benchmark for model accuracy, specificity, and F1 score with respect to macrophage phenotype classification.

Topological Data Analysis: Topological data analysis (TDA) [30] can be broadly defined as a set of data analysis techniques aimed at discovering hidden meaningful structure from high-dimensional data. Persistent homology and the Mapper [30] are two of the most often used TDA approaches. In our current study we explored TDA-mapper to analyze our data. Mapper is an TDA algorithm which employs dimensionality reduction, clustering, and graphical networks for generating an improved visualization and comprehension of higher dimensional data. The working principle of the mapper algorithm is described below:

1. Project the higher dimensional data into low dimension using a filter function (e.g., PCA, Uniform Manifold Approximation and Projection (UMAP))
2. Cover this projection with overlapping hypercubes/intervals with uniform length. 3) Cluster data points within an interval (e.g., Hierarchical clustering, DBSCAN, etc.). 4) Establish a graphical network with vertices representing cluster sets, and edges connecting clusters that share points with some similarity.

TDA was applied to the metabolic datasets from supernatant and cell pellet which contained the 9 samples as rows but only the top metabolites as columns (HMDB ID). For this study, we utilized UMAP and DBSCAN as filtering and clustering algorithms, respectively. Cubicalcover with an overlap fraction of 0.3 was used to create overlapping hypercubes. Giotto-TDA toolbox [https://giotto-ai.github.io/gtda-docs/0.5.1/library.html] was used to develop the mapper network.

## 3. Results

### 3.1 Models of macrophage polarization

Polarization of macrophages from naïve (M0) to anti-tumor M1 or protumor M2 type is mediated by a broad range of biological signals such as cytokines (TNFα, IL-1β, IL-17, TGF-β, IL-13 and IL-4), chemokines (Fig 1A) and extracellular vesicles (EVs, Fig 1B) secreted by the tumor cells. All the terms and the units used in the equations moldeing the interactions are described in Fig 1C. The following equations represent various parameters such as the rate of change in the populations of tumor cells, M0/M1/M2 macrophages, and the quantity of tumor-secreted EVs in the simulated tumor microenvironment.

**Figure 1.**
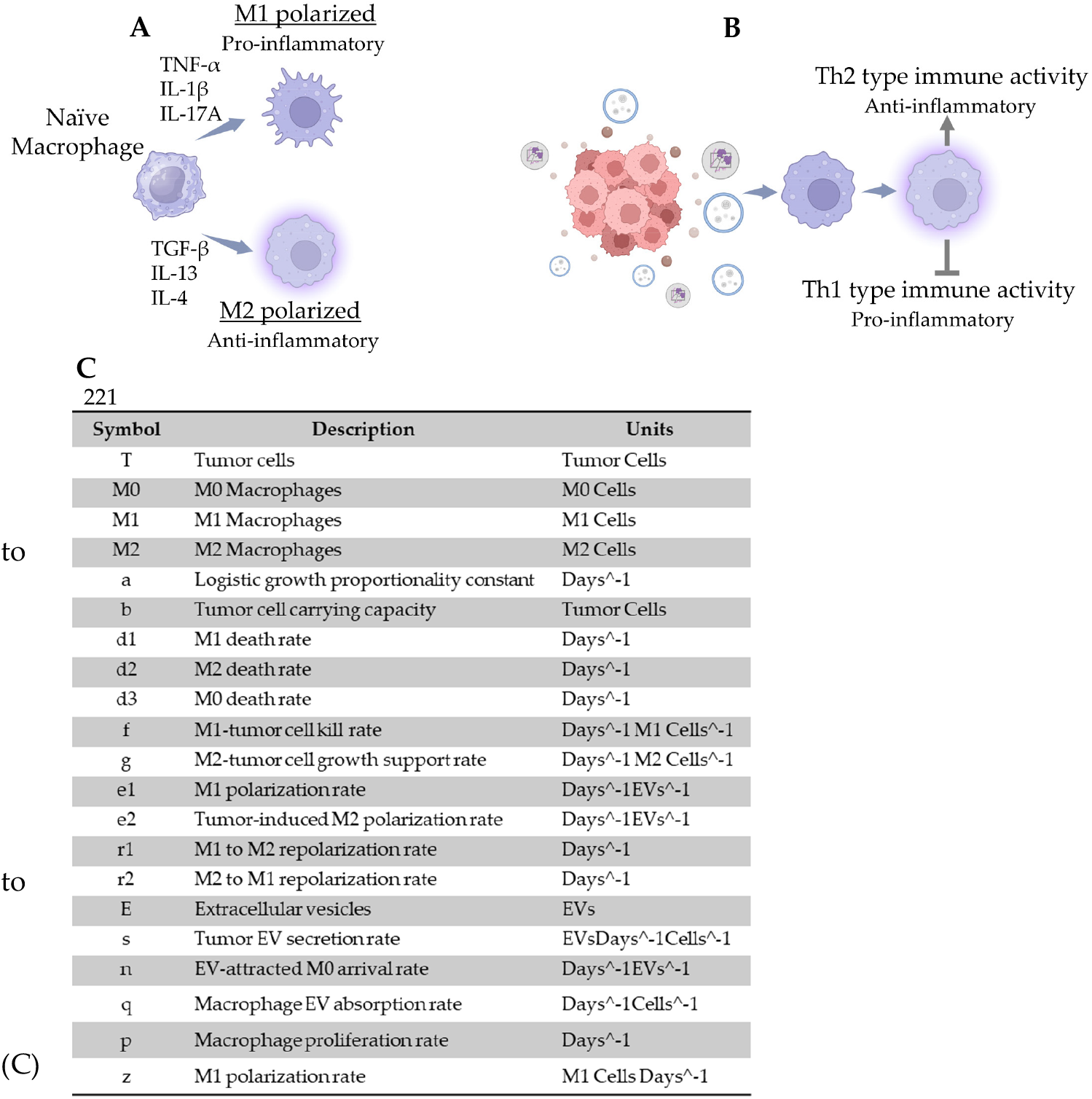
Modeling of the interactions between tumor cells, macrophages, and exosomes. (A) Cytokine-induced polarization of naïve (M0 type) macrophages the M1 and M2 types. TNF-α, IL-1β, and IL-17A are examples of M1-polarizing cytokines; TBF-β, IL-13, and IL-4 are examples of M2-polarizing cytokines. M1 macrophages exhibit pro-inflammatory immune activity, leading to anti-tumor behavior. M2 macrophages exhibit anti-inflammatory activity, leading to pro-tumor growth behavior. B) In the tumor microenvironment, tumor cells release extracellular vesicles that polarize naïve macrophages the M2 subtype. M2 macrophages promote anti-inflammatory Th-2 type immune activity and simultaneously inhibit pro-inflammatory Th-1 type immune activity, supporting tumor cell growth and preventing an effective anti-tumor immune response. All letters/symbols in the ODE system, the corresponding quantities, and the units for each are described.

Equation 1 assumes a logistic growth model for the tumor cell population along with the rates at which M1 macrophages kill tumor cells and M2 macrophages promote tumor cell proliferation.

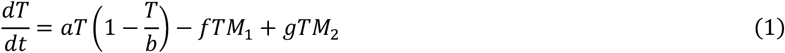

Equation 2 assumes a increasing rate of change of naive (M0) macrophages in the TME, as more immune cells are recruited to the site as time progresses. The rate of change is limited by exosome-dependent polarization to the M1/M2 types. The death rate is a non-factor because once monocytes differentiate into macrophages, the lifespan increases to the range of months. A purely constant M1 polarization term is included to illustrate the impact of M0 polarization to M1 on the tumor cell population.

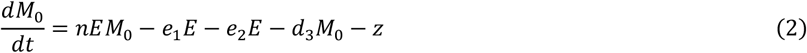

Equations 3 and 4 assume a linearly increasing proliferation rate of M1 macrophages, a polarization rate from M0 based on the number of exosomes, and baseline repolarization rates in either direction. The death rate is also included but assumed to be a non-factor due to the lengthened lifespan after differentiation, similar to the M0 type in equation 2.

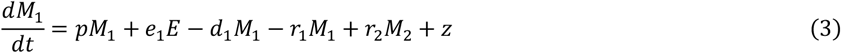

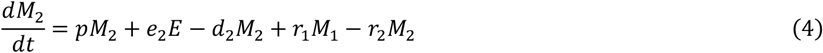

Equation 5 assumes an increase in exosome quantity dependent on the tumor cell population T. The TME exosome quantity decreases due to the nonlinear outflow term that is dependent on E, M0/M1/M2, and constant q as macrophages absorb the exosomes.

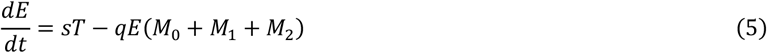

### 3.2 Rate of M1 polarization is the critical parameter for anti-tumor activity

The system of differential equations was solved to obtain solution curves for each population in the system. To demonstrate the importance of polarization towards the M1 type in reducing tumor cell population, the M1 polarization parameter z was swept starting at 0 up to 550 M1 cells per day (Fig 2A-E) and the numerical values described (Fig 2F). Even with this term held constant, there was a reduction in the tumor cell population of up to 7 orders of magnitude while all other parameters remained unchanged (Fig 2A and 2E). This demonstrated the significant effect that even a comparatively smaller M1 polarization rate can have in eliminating tumor cells in the tumor microenvironment.

**Figure 2.**
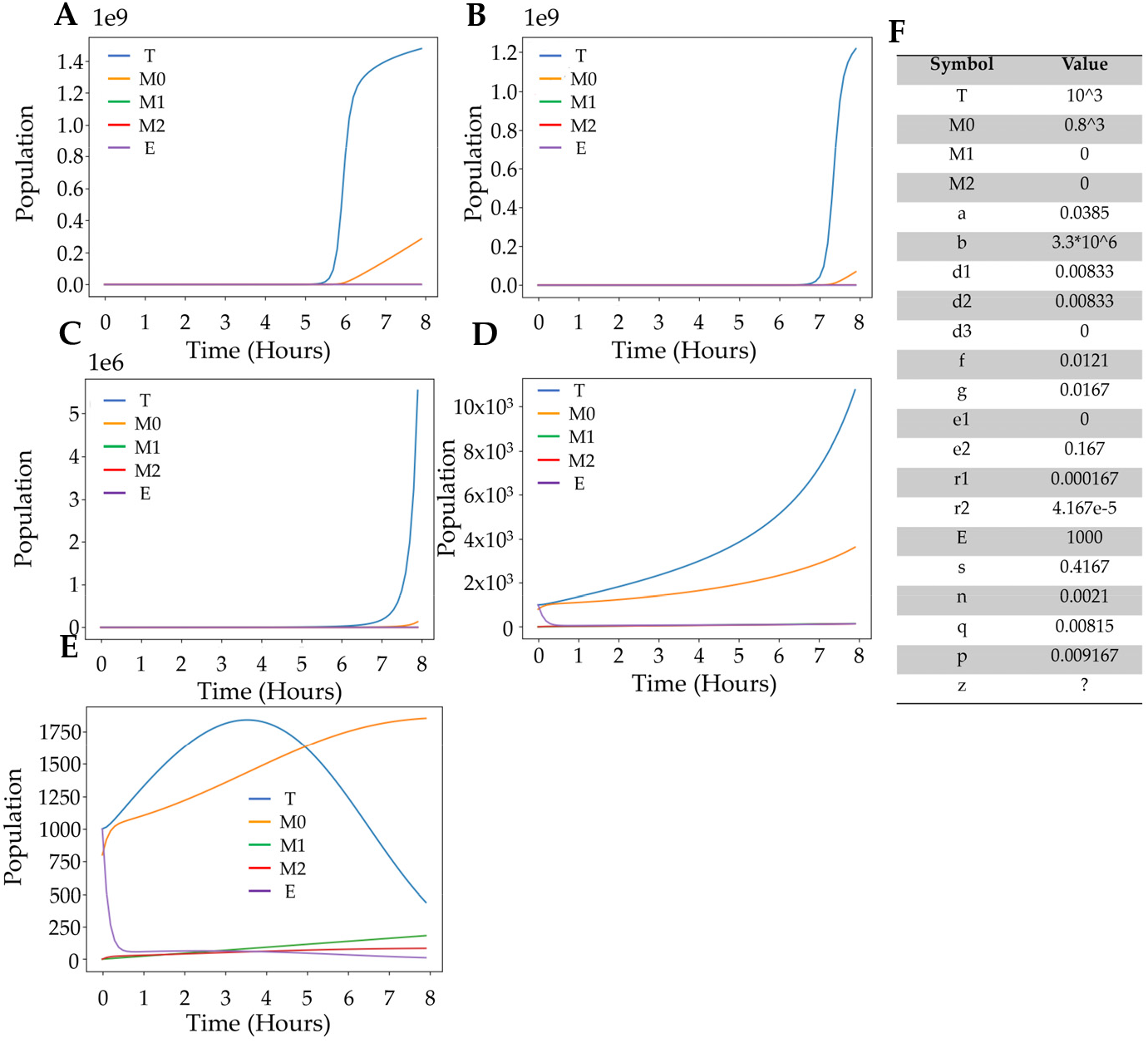
Identification of critical parameter in anti-tumor activity by macrophages. If a negligible amount of macrophages are being polarized to the M1 phenotype represented by z, and no repolarization is occurring, the tumor cell population becomes the greatest in the microenvironment by several orders of magnitude above all non-M0 populations. Keeping all other coefficients constant and sweeping the z parameter 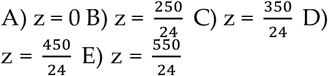 revealed that increasing the rate of M1 polarization mitigates tumor cell population growth. F) The numerical values obtained from the literature used in the simluations are listed.

### 3.3 Identification of metabolites associated with macrophage polarization

Macrophage polarization is accompanied by rapid changes in the metabolism of the cells. To explore the role of metabolites associated with M1/M2 polarization, we performed untargeted mass spectrometric analysis of metabolites extracted from *in vitro* differentiated RAW 246.7 macrophages. In vitro differentiation was accomplished using standard cytokine treatment protocols (Fig 3A). Both the cell superntants and the cell pellets were analyzed by LC/MS to identify secreted and cell associated metabolites associated with polarization.

**Figure 3.**
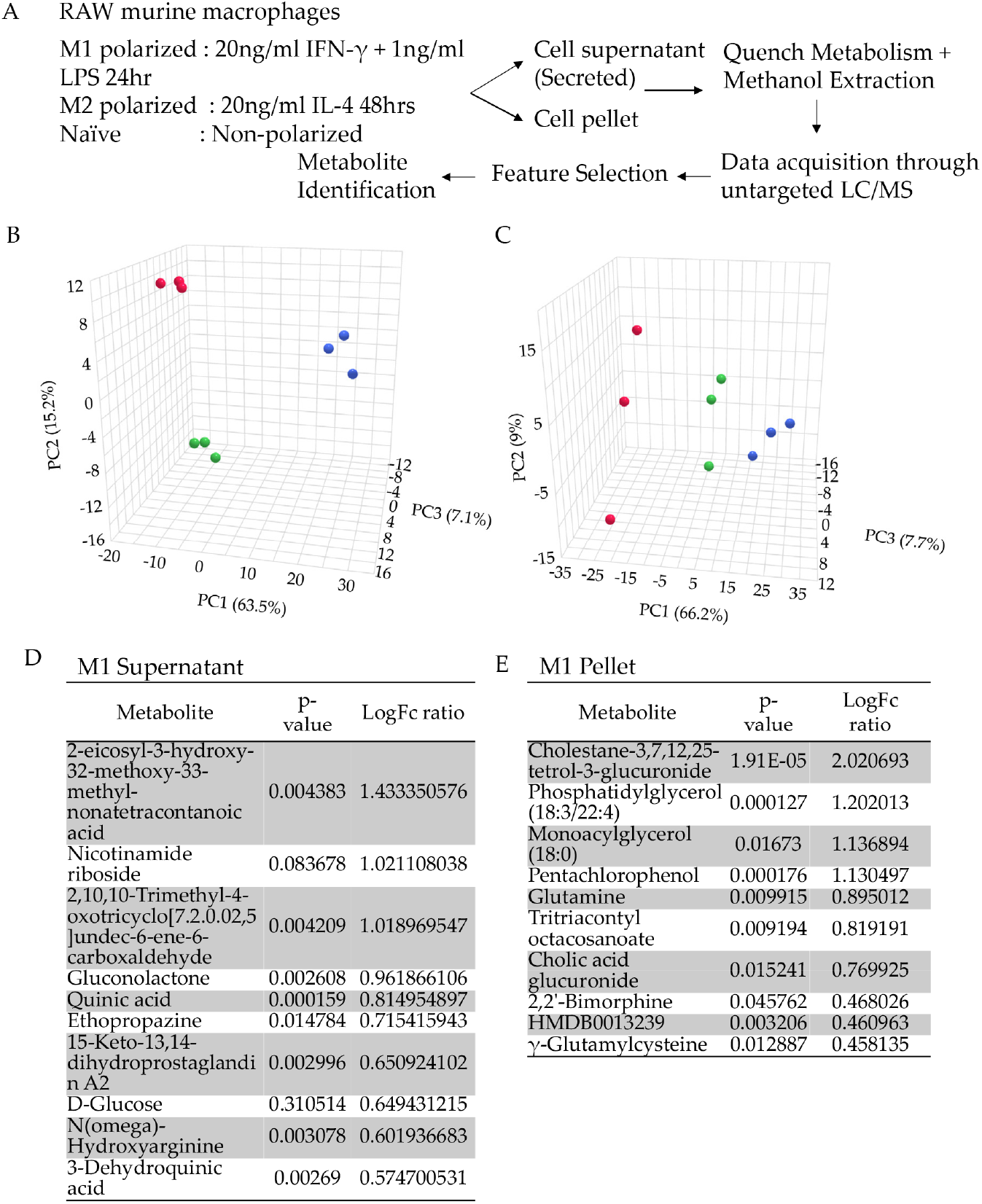
Metabolomic analysis of polarized and naïve macrophages. A) RAW macrophages were polarized by treating with IFN-γ and LPS or IL-4 to obtain M1 or M2 macrophages. Non-polarized macrophages were used as naïve controls. Cell supernatants and pellets were processed to identify metabolites that are either secreted and cell-associated by LC/MS. Principal component analysis on the metabolite abundances in naïve, M1 or M2 polarized macrophages for (B) supernatant and (C) cell pellet showed non-overlapping sample space indicating the clear biological distinction between the groups. The log fold-enrichment of metabolites in M1 macrophages compared to naïve for supernatant (D) and cell pellet (E) samples are shown.

A total of 503 annotated metabolites were identified in the supernatants whereas a total of 663 annotated metaboliates were identified from the cell pellets (Supplementary information). Principal component analysis (PCA) was performed using raw abundance data of each metabolite for M1(red dots), M2 (green dots), and naïve (blue dots) macrophage samples, each with three replicates, in the supernatant (Fig 3B) and cell pellet (Fig 3C) separately. Both analyses produced score plots with three non-overlapping clusters, indicating that the metabolic profile of each phenotype was biologically distinct. The top 10 annotated metabolites enriched in the supernants (Fig 3D) and cell pellets (Fig 3E) of M1 polarized macrophages sorted by fold change compared to naïve (M0) macropahges (LogFc ratio) are shown. Distinct molecules were found to be enriched in supernatants and cell pellets. The complete list of metabolites detected with m/z ratio, retention time, raw and normalized abundances are shown in the supplmentary information.

### 3.4 Ensemble methods identify metabolites associated with macrophage polarization

Selecting metabolites based on the magnitude of the fold-change ratio or its logarithm to identify the important metabolites fails to take into account the differences in scaling, as it does not directly compare the raw abundances found from LC/MS data. To overcome this limitation, a binary classifier for M1 macrophages was built using the random forest classifier model to determine the metabolite of the highest importance in determining the macrophage phenotype, using both the supernatant and cell pellet data separately (Fig 4A). Chi-square, logistic regression, and ANOVA F-value procedures were used as feature selection methods to narrow down the possible metabolites to a list of only the most determinant ones. As the model was repeatedly trained, the most important metabolites were continuously tracked. In the end, the metabolites with the highest number of selection instances were retained, revealing the most important metabolites overall across all three methods.

**Figure 4.**
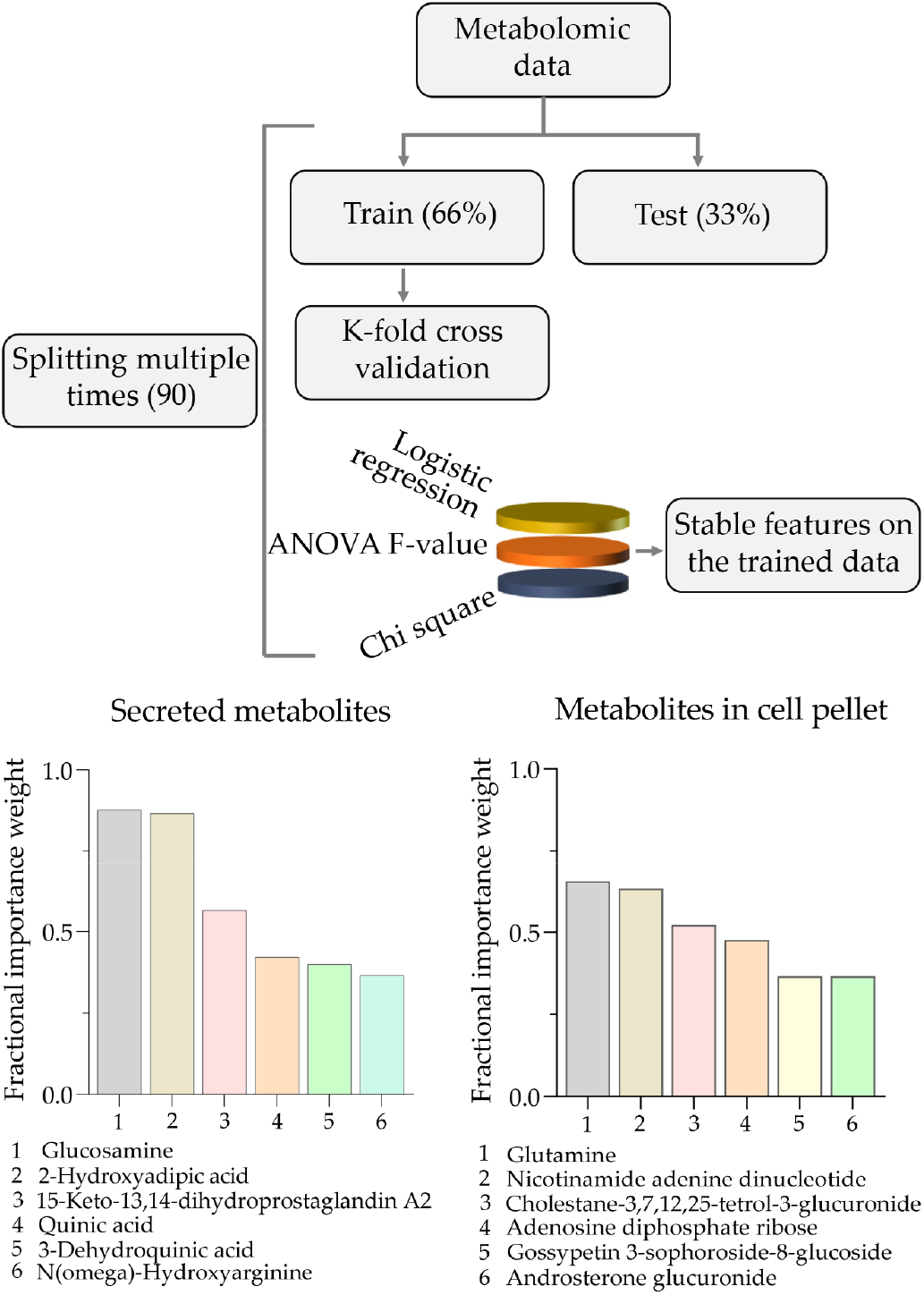
Selection of metabolites based on ensemble ML method: A) The metabolomic data from supernatant and cell pellets were split into training and test sets for cross validation. Stable features were selected based on three feature selection methods. Metabolites with highest predictive weight of M1 polarization from supernatant (B) and cell pellet (C) are shown.

In contrast to the fold enrichment methods, our model selects the most important features in each iteration based on the chi-square, logistic regression, and ANOVA F-value statistics. The model uses the raw abundance as input and does not consider the relationships between only M1 and naive macrophage abundance, avoiding the problem of scaling with the raw values. The applied feature selection methods were chosen due to the fact that they are better suited for datasets with few samples, such as the metabolomics data in this analysis.

Results showed that mainly metabolites associated with glycolytic and lipid metabolism were enriched in both the supernatant (Fig 4B) and cell pellets (Fig 4C). Glucosamine and 2-hydroxyadipic acid were selected among the most important features in predicting M1 polarization 87% of the time in the secreted metabolite set (Fig 4B) whereas the presence of glutamine was the top selected metabolite (65.5% of the time) to predict M1 polarization in the cell associated metabolite dataset (Fig 4C). Results from our model showed non-overlapping sets of top metabolites when compared with the fold enrichment ranks (Fig 3D-E) indicating the power of the random forest classifer in the analysis of metabolomic datasets.

**Table 1.** Classification performance across methods, using all or selected features. To assess the classification power of the selected metabolites, the random forest classifier was compared to logistic regression and support vector machine (SVM) classifiers. Accuracy, precision, and F1 score were found to be 3-4% lower when using all features for the random forest classifier compared to using only the selected features. Similarly, The SVM classifier had 5% lower accuracy and 14% lower recall and F1 score when using all features compared to only those that were selected. All methods showed 100% accuracy, precision, recall, and F1 score when only the selected features were used, indicating the importance of the top metabolites in accurately characterizing the phenotype of the macrophages.

| Classification Method | All Features |  |  | Selected Features |  |  |
| --- | --- | --- | --- | --- | --- | --- |
|  | RF | Logistic Regression | SVM | RF | Logistic Regression | SVM |
| Accuracy | 0.97 | 1.00 | 0.95 | 1.00 | 1.00 | 1.00 |
| Precision | 0.96 | 1.00 | 1.0 | 1.00 | 1.00 | 1.00 |
| Recall | 1.00 | 1.00 | 0.86 | 1.00 | 1.00 | 1.00 |
| F1 Score | 0.97 | 1.00 | 0.86 | 1.00 | 1.00 | 1.00 |

### 3.5 Topological data analysis confirms the results from ML analysis

To confirm the validity of the findings from ensemble ML analysis, we performed topological data analysis (TDA) on the selected features. The top 6 features from the supernatant and cell pellet analysis were utilized for the TDA mapper visualization. The Mapper graphs illustrated interesting formation of sample clusters by the topological structure. Two separated clusters were formed: all samples from the M1 class (cluster with red node) and all samples from non-M1 class (cluster with blue node), which showed that combination of these six features differentiate the phenotypes. TDA mapper graphs for the selected features from supernatant and pellets are shown in figure 5. The clustering patterns for the pellets differed from that of supernatant presumably due to the presence of outliers.

**Figure 5.**
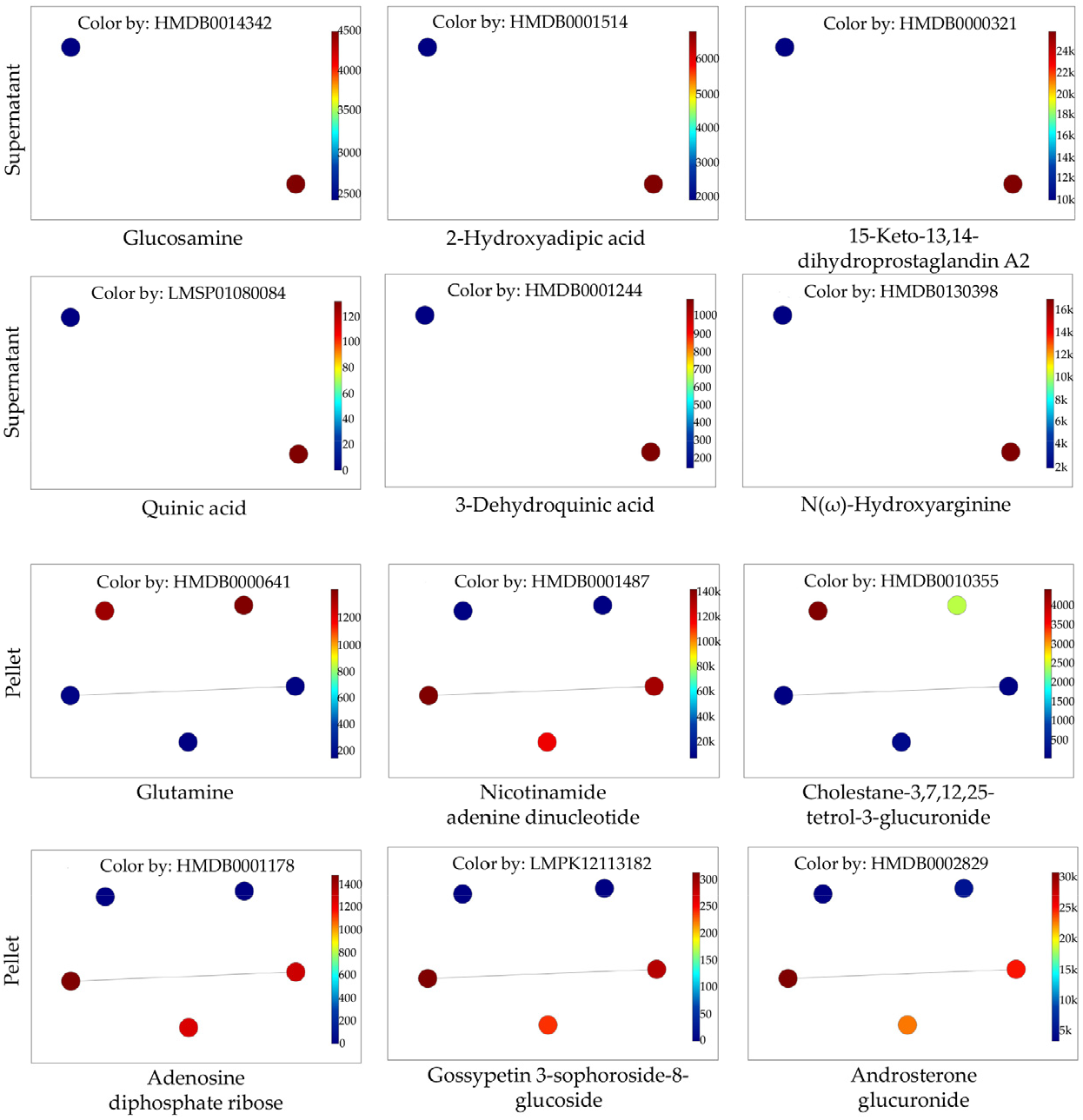
Visualization of features by TDA mapper. Selection of metabolites was based on ensemble ML method. A graphical network with vertices representing cluster sets, and edges connecting clusters that share points with some similarity was created to visualize the features. Metabolites from the supernatant (top panels) and the pellets (bottom panels) predicting the M1 (red dots) and non-M1 (blue dots) phenotypes are shown.

## 4. Discussion

In this manuscript we designed a system of ordinary differential equations described by equations (1) – (5) to model the populations of tumor cells, M0/M1/M2 macrophages, and the quantity of EVs in a TME in a predator-prey-style simulation. Tumor cells represent the prey and M1 macrophages represent the predators, and the populations of M2 macrophages and EVs are there to support tumor cell growth. M0 macrophages being the precursors, supply both the M1 and M2 populations. EVs may also be considered prey of the macrophages, as they are being introduced into the simulation from tumor cells and removed via absorption by the macrophages. The population values were scaled down by a factor of 10^3^ from typical experimental values to reduce computational time. Additionally, EVs may act as decoys such that the macrophages or other immune cells attack the EVs and are diverted from tumor killing. In this scenario, the EV population would compete with the tumor cells as prey for the macrophages, and would be a positive variable that would promote the cancer. A limitation of our model is that since our equations represent a predator-prey simulation, only the populations are being modeled without a spatial dimension.

To ensure robustness, the parameters used in the ODEs described by equations (1) – (5) were obtained from experiments or similar models in the literature and also scaled down by a factor of 10^3^ if dimensionally necessary. The initial M0 population was based on *in vitro* experiments in 6-well plates seeded with THP-1 monocytes followed by polarization [31]. The initial tumor cell population was 200 cells higher to represent the tumor cells as the largest population in the initial TME. The initial M1 and M2 populations were both set at 0 as minimal polarization would occur as M0 macrophages first arrive at the TME. The initial EV quantity was assumed to be the same as the tumor cell count since the model was taking into account the effects of EVs secreted in response to the recruitment of macrophages. The rate of EV secretion by tumor cells was obtained from previous studies on tumor EVs [32, 33]. The natural death rate of M0 macrophages (represented by *d3*) was assumed to be 0, as once monocytes differentiate into naive macrophages the lifespan lengthens to a range of months and years [34]. The rate of EV absorption by macrophages (*q*) was obtained from experiments on EV uptake by RAW 246.7 macrophages [35]. The rate of M1 polarization (z) was set to 0, as all tumor cell EVs were assumed to induce polarization towards the tumor-supportive M2 type. The EV-attracted M0 arrival rate (*n*) was set to 0.05 EVs^-1Days^-1. Since this is two orders of magnitude less than the tumor cell EV secretion rate, the assumed value is thus reasonable for the simulation. All other parameters (a, b, d1, d2, f, g, e2, r1, r2, q, p) described in the ODE model were based on sources in the literature [36].

Sweeping the *z* parameter for polarization toward the M1 type increased the M1 population and substantially reduced the tumor cell population as the parameter increased in value. Over the full duration of the simulation, the M1 polarization parameter is relatively small in magnitude, as it is the only constant term in the system. Thus, the model demonstrates that even a constant, comparatively small rate of M1 polarization to increase the M1 population is highly effective at killing the tumor cells, suggesting pharmacological intervention targeting M1 polarization could be effective. Since the metabolic activity of macrophages determines their function, promoting M1-characteristic metabolism can polarize towards the tumoricidal phenotype.

Previous studies identified metabolic pathways that could be targeted to induce a specific phenotype [7, 9-11]. Towards identifying metabolites associated with M1 or M2 phenotype, we analyzed metabolomic data from an untargeted LC/MS experiment. Conventional methods based on enrichment ratios may not reveal the *bona fide* biomarkers of macrophage polarization. Therefore, we employed ensemble machine learning analysis that takes into consideration the broad range in the experimentally observed data of the metabolite abundances, and discriminates between macrophage phenotypes. Our methods identify the most important metabolites in classifying a macrophage as M1 based on the metabolic profile. We considered both cell-associated and secreted (culture supernatant) metabolites in our analyses with the assumption that the metabolites presumably secreted as cargo in macrophage EVs may also affect the polarization. A random forest classifier was implemented due to the small number of samples and high dimensionality of the data. Random forest is considered an optimal ML method for metabolite selection based on classification using LC-MS data [37]. RFs are composed of many decision trees trained on a subset of features, which reduces the risk of overfitting with the high-feature dataset compared to a single decision tree. Additionally, ML based methods are better suited when constructing a model to analyze data obtained from stochastic processes and to infer dynamical changes. For example, data on the movement dynamics of antelopes utilized models of diffusion to predict herd movement [38].

Among the top five selections in the supernatant data, we found metabolites involved in glycolytic metabolism regulation and lipid metabolism as well as chlorogenic acids. For instance, glucosamine is known to cause a reduction in glycolytic metabolism via the inhibition of glyceraldehyde-3-P dehydrogenase and lactate dehydrogenase [39], and therefore causes suppression of M1 gene expression, making it critical for polarization [40]. Chlorogenic acids, which can be derived from the identified metabolites quinic acid and 3-Dehydroquinic acid, have been shown to promote M1 polarization that ultimately limits tumor cell growth [41]. 15-Keto-13,14-dihydroprostaglandin A2 (a prostanoid) and 2-hydroxyadipic acid (a fatty acid) were also identified to be associated with macrophage polarization. Experimental research confirms that lipid metabolism, specifically fatty acid oxidation (FAO), is higher in M2 macrophages compared to M1 macrophages [42].

Similarly, the top five selections in the cell pellet data (Fig 4C) comprise metabolites involved in glycolytic metabolism, aerobic metabolic pathways (OXPHOS and TCA cycle), or lipid metabolism. Glutamine inhibition has been shown to increase glucose flux through glycolysis, promoting M1 polarization for an anti-metastatic effect [43]. Nicotinamide adenine dinucleotide (NAD) and adenosine diphosphate ribose (ADPR) are substrates in the ATP-generating metabolic pathways characteristic of M2 macrophages. Cholestane-3,7,12,25-tetrol-3-glucuronide and gossypetin 3-sophoroside-8-glucoside are also lipids and polyketides that may be involved in lipid metabolism that influences macrophages phenotype as discussed above.

The top metabolites are selected based on feature selection across 90 total training iterations, 30 for each feature selection method (chi-square, logistic regression, and ANOVA F-value). As our dataset is smaller, we chose simpler feature selection algorithms. Ensemble feature selection would mitigate bias and generate stable biomarkers. Stratified K-fold cross-validation was used so 6 samples were used for training and 3 for testing in each iteration. Nested cross-validation was conducted using the leave-one-out (LOO) approach to avoid overfitting due to information leakage in simple cross-validation due to the small sample number. Due to class imbalance in M1 vs. non-M1 samples, SMOTE was applied with one nearest neighbor on the outer loop training set, so the classes were balanced before feature selection.

Previous studies that used non-ML methods identified itaconate, succinate, and citrate are known to accumulate in M1 macrophages and therefore play an important role in determining the phenotype [44, 45]. The fitness of our ensemble methods for the identification of biomarkers of macrophage polarization is revealed by the subset of metabolites identified by the model are similar in their biochemical properties in glycolytic, aerobic and lipid metabolism as shown in previous studies using non-ML methods. Additionally, studies on human macrophages polarized *in vitro* and analyzed using ML methods such as random forest for classification identified metabolites belonging to fattyacid metabolism as the discriminating features between M1 and M2 macrophages, similar to our results using murine macrophages suggesting that the methods produced comparable results [46].

In order to verify if the top metabolites identified in the model have a polarizing effect, a biological experiment must be conducted. To test a certain metabolite, a sample of M0 macrophages must be treated with an agonist/inhibitor of the metabolic pathway that produces the metabolite. If M1 polarization is indeed observed, the macrophages may be placed in a simulated TME with tumor cells. If a decline in tumor cell population due to tumoricidal activity by M1 macrophages is observed, the selected metabolite can be established as a viable target for anti-tumor M1 polarization. Such approaches may be combined with the anti-cancer chemotherapies to enhance the efficacy of tumor killing.

## Supporting information

Supplemental 1 for Figure 4

Supplemental 2 for Figure 4

Supplemental 1 for Figure 3

Supplemental 2 for Figure 3

## Supplementary Materials

Raw metabolite abundances from the mass spectrometric analysis of cell pellet and cell supernatants is provided as supplementary information.

## Author Contributions

“Conceptualization, A.G., R.P., S.K., and P.S.; methodology, A.G., R.P., S.K., P.S.; software, R.P., S.K..; validation, A.G., R.P.; formal analysis, A.G., S.K. R.P., P.S.; investigation, A.G., M.H., L.L.; resources, M.H., L.L.; data curation, A.G., S.K., R.P.; writing—original draft preparation, A.G.; writing-review and editing, A.G., R.P. S.K., L.L, P.S.; visualization, A.G.; supervision, R.P., S.K., P.S.; project administration, L.L., P.S. All authors have read and agreed to the published version of the manuscript.”

## Data Availability Statement

All the mass-spectrometry data used in the metabolomics analysis is submitted as supplementary information. The code used in the analysis will be made available upon request.

## Acknowledgments

[AG] is a recipient of summer internship opportunities through the ‘Aspiring Scientists Summer Internship Program’ at George Mason University in the summer of 2023 [under Prof Lance Liotta] and 2024 [under Prof. Padmanabhan Seshaiyer].

## Disclaimer/Publisher’s Note

The statements, opinions and data contained in all publications are solely those of the individual author(s) and contributor(s) and not of MDPI and/or the editor(s). MDPI and/or the editor(s) disclaim responsibility for any injury to people or property resulting from any ideas, methods, instructions or products referred to in the content.

